# Intranasal immunization with live-attenuated RSV-vectored SARS-CoV-2 vaccines elicits antigen-specific systemic and mucosal immunity and protects against viral challenge and natural infection

**DOI:** 10.64898/2026.03.30.715311

**Authors:** Davide Botta, Michael D. Schultz, Aaron Silva-Sanchez, Davies Kalange, Jobaida Akther, Fen Zhou, Jennifer L. Tipper, Guang Yang, Levi T. Schaefers, Courtney A. Barkley, Shihong Qiu, Jeremy B. Foote, Mariana F. Tioni, Christopher M. Weiss, Shannon I. Phan, Todd J. Green, Sixto M. Leal, Kevin S. Harrod, Rodney G. King, Martin L. Moore, Troy D. Randall, Roderick S. Tang, Frances E. Lund

**Affiliations:** Department of Microbiology, The University of Alabama at Birmingham, Birmingham, AL, USA; Immunology Institute, The University of Alabama at Birmingham, Birmingham, AL, USA; Department of Medicine, Division of Clinical Immunology and Rheumatology, The University of Alabama at Birmingham, Birmingham, AL, USA; Department of Anesthesiology and Perioperative Medicine, The University of Alabama at Birmingham, Birmingham, AL, USA; Department of Pathology, Division of Laboratory Medicine, The University of Alabama at Birmingham, Birmingham, AL, USA; Meissa Vaccines Inc, Redwood City, CA, USA

**Keywords:** SARS-CoV-2, Severe acute respiratory syndrome coronavirus 2 RSV, Respiratory syncytial virus, CBA Cytometric bead array, mLN Mediastinal lymph node, URT Upper respiratory tract, LRT Lower respiratory tract

## Abstract

The emergence of new SARS-CoV-2 variants and breakthrough infections underscores the need for next-generation vaccines capable of protecting from natural infection and/or preventing virus transmission to others. Intranasal vaccination offers a promising approach by eliciting local immune responses in the nasal mucosa, the primary site of infection and reservoir for transmissible virus. We evaluated two live-attenuated, respiratory syncytial virus vectored vaccines in which the RSV F and G surface glycoproteins were replaced with a chimeric SARS-CoV-2 Spike protein from either the ancestral USA/WA-1/2020 strain (MV-014-212) or the Delta variant (MV-014-212-delta). A single intranasal dose of either vaccine elicited systemic and mucosal immunity in K18-hACE2 mice, including serum neutralizing antibodies, Spike-specific memory B cells and plasmablasts, and Spike-specific CD8⁺ lung-resident memory T cells. Although MV-014-212-delta vaccination provided the best protection against Delta variant virus challenge, both vaccines decreased viral loads in nasal discharge, lung and brain, and reduced weight loss and mortality. In naturally acquired infection studies, vaccinated hamsters exposed to infected cagemates exhibited minimal weight loss, limited viral replication within the nasal mucosa, and attenuated lung pathology. Therefore, intranasal RSV-vectored vaccines can elicit broad protective respiratory immunity, suggesting that this platform could be leveraged for other respiratory pathogens.

## 1. Introduction

Severe acute respiratory syndrome coronavirus 2 (SARS-CoV-2) is the etiologic agent responsible for the 2019 coronavirus disease (COVID-19) pandemic. According to the World Health Organization (WHO) Coronavirus Disease Dashboard [1], more than 778 million cases have been reported globally, including over 103 million in the United States as of February 2025. Transmission occurs primarily through exposure to virus-laden respiratory droplets, aerosols, and contaminated surfaces [2, 3]. Infected individuals may experience a spectrum of clinical outcomes, ranging from asymptomatic to severe and life-threatening disease. The high transmissibility of SARS-CoV-2 continues to pose significant challenges for global containment, underscoring the need for vaccines capable of protecting vaccine recipients from naturally (community)-acquired infections and able to limit virus transmission between the vaccinated individual and others, particularly those with weakened immune systems.

While the gold standard for vaccines is sterilizing immunity, which is defined as the complete clearance of a pathogen at or near the site of entry [4] many vaccines are unable to reach this high bar. However, even non-sterilizing immunity can decrease pathogen replication and pathogen load, which ultimately leads to reduced morbidity and mortality and a shorter time-period in which transmission of the pathogen from the vaccinated subject to a non-immune bystander can occur [4]. Both non-sterilizing and sterilizing immunity can benefit from a strong systemic immune response and from local immunity that is specifically focused on the site of infection [4, 5]. Although local and systemic immune responses readily engage the innate, cellular and humoral arms of the immune system, the most effective vaccines, at least to date, work by eliciting durable, antigen-specific antibody responses through T cell-dependent affinity maturation [4, 5]. For pathogens that target mucosal surfaces, secretory IgA serves as the principal antibody isotype [5] and it is appreciated that elevated mucosal IgA titers are associated with resolution of respiratory virus infections [6, 7], including SARS-CoV-2 [8, 9]. Moreover, memory T and B cells residing in the upper respiratory tract (URT) and the lower respiratory tract (LRT) can also support more rapid pathogen clearance following infections with pathogens that target the respiratory tract [6, 10–12]. Thus, the mucosal immune response can play an essential role in containing respiratory virus infections.

Despite the clear role for the mucosal immune response in resolving respiratory tract infections, vaccination strategies typically rely on systemic delivery of antigen. Indeed, as of 2025, only five mucosal vaccines for SARS-CoV-2 have received approval for use in humans, and all are administered outside the United States [13–18]. Instead, almost all FDA-approved vaccines in the US are administered via the intramuscular route [19], which elicits rapid antigen uptake and processing within highly vascularized muscle tissue [20]. While these intramuscular vaccines can induce robust systemic IgG responses [21] and effector and memory cells that reside in secondary lymphoid tissues [22], the mucosal response elicited by intramuscular vaccines is limited, and these vaccines fail to generate durable, antigen-specific IgA responses [23–26] and mucosal resident memory cells [27]. This lack of local immunity can preclude establishment of sterilizing immunity to respiratory pathogens [23], particularly when the pathogen can rapidly mutate to escape the systemic neutralizing antibody response. Thus, individuals vaccinated via the intramuscular route may become infected through exposure to infected individuals in their environment, and while the vaccinated individual will likely be protected from severe disease manifestations [26], these individuals may remain capable of transmitting the virus to others [5, 26]. Moreover, viral replication in asymptomatic, vaccinated hosts may also increase the potential for the emergence of vaccine-resistant variants [5]. Therefore, development of vaccines capable of inducing both systemic immunity and mucosal immunity at the site of pathogen entry is of critical importance.

The mucosal surface of the lung and nasal cavity, which is lined with permeable epithelium and enriched with resident immune cells [28], serves as the primary entry point of respiratory pathogens like SARS-CoV-2 [29]. Preclinical studies using different vaccine platforms, including replication deficient adenovirus vectors and various virus-like particles, to deliver coronavirus antigens via the intranasal route have demonstrated durable systemic and mucosal immune responses in mice and hamsters [30–34], as well as humans [35–37]. Consistent with these results, we previously showed that intranasal (i.n.) delivery of a live-attenuated, respiratory syncytial virus (RSV)-vectored vaccine, in which the RSV F and G surface glycoproteins were replaced with a chimeric SARS-CoV-2 Spike protein, was sufficient to elicit robust T cell-mediated antibody response in non-human primates following a single i.n. dose of the vaccine [38]. Given that i.n. delivery of the RSV-vectored vaccine platform induced strong immunity, we hypothesized that this vaccine would effectively protect animals from naturally acquired infection. To test this, we evaluated two i.n. administered RSV-vectored vaccine candidates engineered to express the SARS-CoV-2 Spike protein cloned from either the ancestral USA/WA-1/2020 strain (MV-014-212) or the Delta (B.1.617.2) variant (MV-014-212-delta). Following a single i.n. dose of the vaccines, we observed Spike-specific cellular, humoral, and mucosal immune responses in K18-hACE2 transgenic mice. Furthermore, vaccinated animals exhibited markedly reduced viral burden, morbidity, and mortality following live SARS-CoV-2 challenge. Importantly, hamsters immunized with the RSV-vectored vaccines were more protected from naturally acquired infection with SARS-CoV-2 when compared to sham-vaccinated animals. Therefore, the attenuated RSV-vector vaccine platform can be used to drive protective systemic and local mucosal immunity that can limit community acquired infection.

## 2. Results

### 2.1 A single i.n. administration of MV-014-212 or MV-014-212-delta induces systemic vaccine-specific neutralizing antibody responses in mice

To evaluate the immunogenicity of MV-014-212 and MV-014-212-delta vaccines, K18-hACE2 transgenic mice, which express the ACE2 receptor under the control of the K18 promoter [39], were administered a single i.n. dose of the vaccines. As expected, sham-vaccinated mice that received a single i.n. administration of Vero cell lysate failed to develop IgM (**Figure S1a and S1b**) or IgG (**Figure 1a and 1b**) antibody responses specific for the full ectodomain of SARS-CoV-2 Spike or the Spike Receptor Binding Domain (RBD). In contrast, mice vaccinated i.n. with either MV-014-212 or MV-014-212-delta seroconverted, and exhibited early but transient IgM responses specific for Spike and RBD (**Figure S1a-b**) as well as elevated Spike- and RBD-specific IgG responses that were detected as early as day 10 post vaccination and maintained at high levels for at least 5 weeks (**Figures 1a and 1b**). To address the neutralizing capacity of the antibodies elicited by vaccination, we evaluated whether the antibodies were sufficient to block infection of hACE2-expressing cells with a SARS-CoV-2 pseudovirus. We found that day 30 sera from vaccinated animals, but not from sham-vaccinated animals, effectively inhibited the pseudovirus infection *in vitro* with comparable neutralizing activity (**Figure 1c**). Therefore, both MV-014-212 and MV-014-212-delta elicit systemic humoral immunity that is endowed with neutralizing activity.

**Figure 1.**
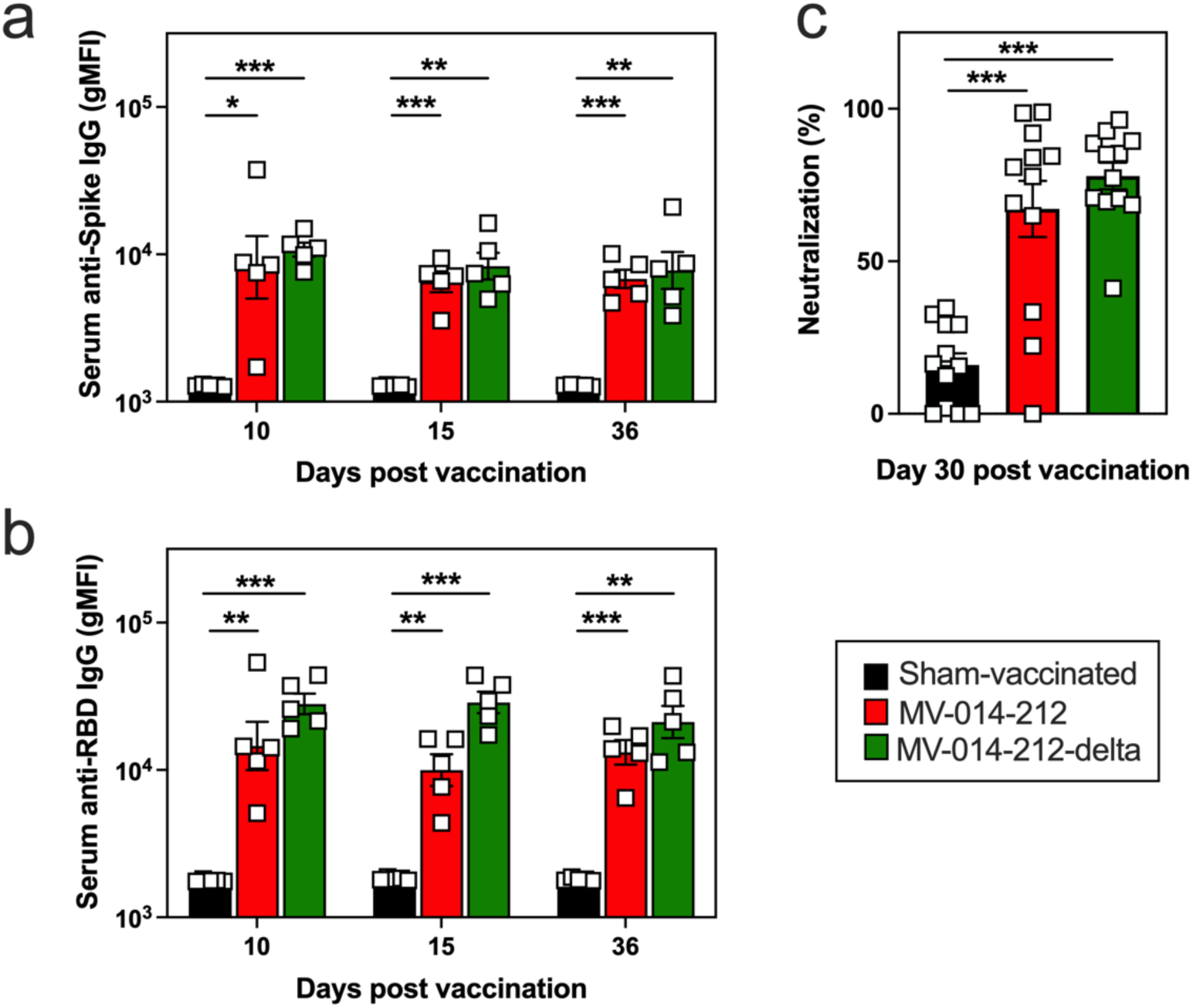
Systemic and neutralizing antigen-specific IgG responses in mice following a single i.n. administration of MV-014-212 or MV-014-212-delta vaccine. Spike-specific (**a**) and RBD-specific (**b**) IgG antibodies in sera from K18-hACE2 mice at days 10, 15 and 36 post vaccination with Vero cell lysate (sham-vaccinated; black), MV-014-212 (red) or MV-014-212-delta (green). Data plotted as log-transformed geometric mean fluorescence intensity (gMFI). (**c**) Normalized percentage neutralization values of sera (diluted 1:100) from K18-hACE2 mice at day 30 post immunization with sham-vaccine, MV-014-212, or MV-014-212-delta. Normalization was calculated by setting control wells containing virus and cells without sera (PBS only) as 0% neutralization. Individual animals (n=5/group for **a-b** and n=12/group for **c**) shown as symbols and group means shown as bars. *P<0.05, **P<0.01, ***P<0.001. P values determined by one-way ANOVA followed by Dunnett’s multiple comparison test. Variability expressed as standard error (SE).

### 2.2 A single i.n. administration of MV-014-212 or MV-014-212-delta elicits vaccine-specific B and CD8⁺ T cell responses in mice

To profile antigen-specific B cell responses, fluorochrome-labeled tetramers of recombinant SARS-CoV-2 Spike trimers were generated to identify Spike-specific germinal center (GC) B cells, plasmablasts, and memory (Mem) B cells (**Figure S2**) within the lung-draining mediastinal lymph node (mLN). Both vaccines induced a robust Spike-specific GC B cell response in the mLN at day 10 post vaccination that was absent in sham-vaccinated mice (**Figures 2a**). Similarly, vaccinated mice developed Spike-specific plasmablasts (**Figure 2b**) and memory B cells (**Figure 2c**) in the mLN, consistent with antigen-specific B cell activation and differentiation.

**Figure 2.**
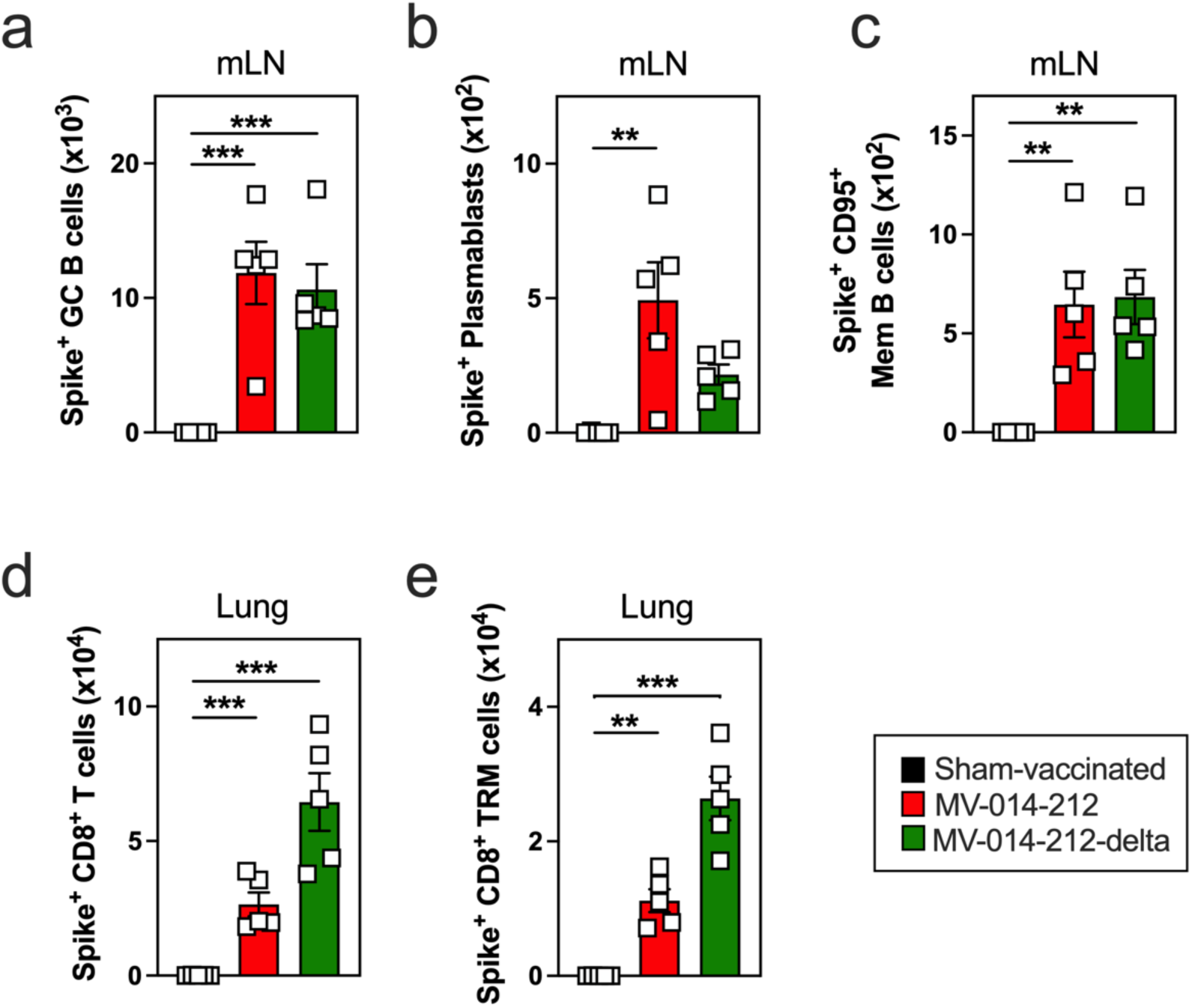
Antigen-specific B and CD8^+^ T cell responses in mice following a single i.n. administration of MV-014-212 or MV-014-212-delta vaccine. Number of Spike-specific germinal center (GC) B cells (**a**), plasmablasts (**b**) and CD95^+^ memory (Mem) B cells (**c**) in the mediastinal lymph node (mLN), and the number of Spike-specific CD8^+^ T cells (**d**) and CD8^+^ tissue-resident memory (TRM) cells (**e**) in the lung from K18-hACE2 mice at day 10 post vaccination with Vero cell lysate (sham-vaccinated; black), MV-014-212 (red) or MV-014-212-delta (green). Individual animals shown as symbols (n=5/group) and group means shown as bars. **P<0.01, ***P<0.001. P values determined by one-way ANOVA followed by Dunnett’s multiple comparison test. Variability expressed as SE.

To assess local cellular immunity, we used Spike peptide loaded Class I tetramers to quantify Spike-specific CD8⁺ T cells in the lung (**Figure S3**) and identify lung-resident memory T (T_RM_) cells – cells that are known to mediate protective responses to SARS-CoV-2 infection [40–42]. Mice vaccinated i.n. using the RSV-vectored platform exhibited significant numbers of Spike-specific CD8⁺ T cells in the lung (**Figure 2d**), and approximately half of these cells co-expressed the canonical markers of tissue residency, namely CD69 and CD103 (**Figure 2e** **and Figure S3**). Therefore, both vaccines can drive robust B and T cell activation and differentiation, and support the formation of antigen-experienced CD8⁺ T_RM_ cells that are known to provide local immune protection against SARS-CoV-2.

### 2.3 A single i.n. administration of MV-014-212 or MV-014-212-delta protects mice against SARS-CoV-2 challenge and limits viral shedding

To determine whether the RSV-vectored vaccines elicited a protective immune response, we challenged the i.n. vaccinated/sham-vaccinated K18-hACE2 mice on day 28 post-vaccination with a lethal dose (2 × 10⁴ PFU) of SARS-CoV-2 Delta variant. As expected, sham-vaccinated mice exhibited severe morbidity, losing up to 20% body weight by day 8 (**Figure 3a**), with 85% mortality by day 9 (**Figure 3b**). In contrast, mice vaccinated with either MV-014-212 or MV-014-212-delta displayed minimal to no weight loss (**Figure 3a**) and markedly improved survival (**Figure 3b**). Although MV-014-212-delta vaccinated mice were fully protected against Delta-induced lethality (**Figure 3b**), MV-014-212 conferred substantial but incomplete protection against the Delta variant challenge virus, with 30% mortality observed at day 9.

**Figure 3.**
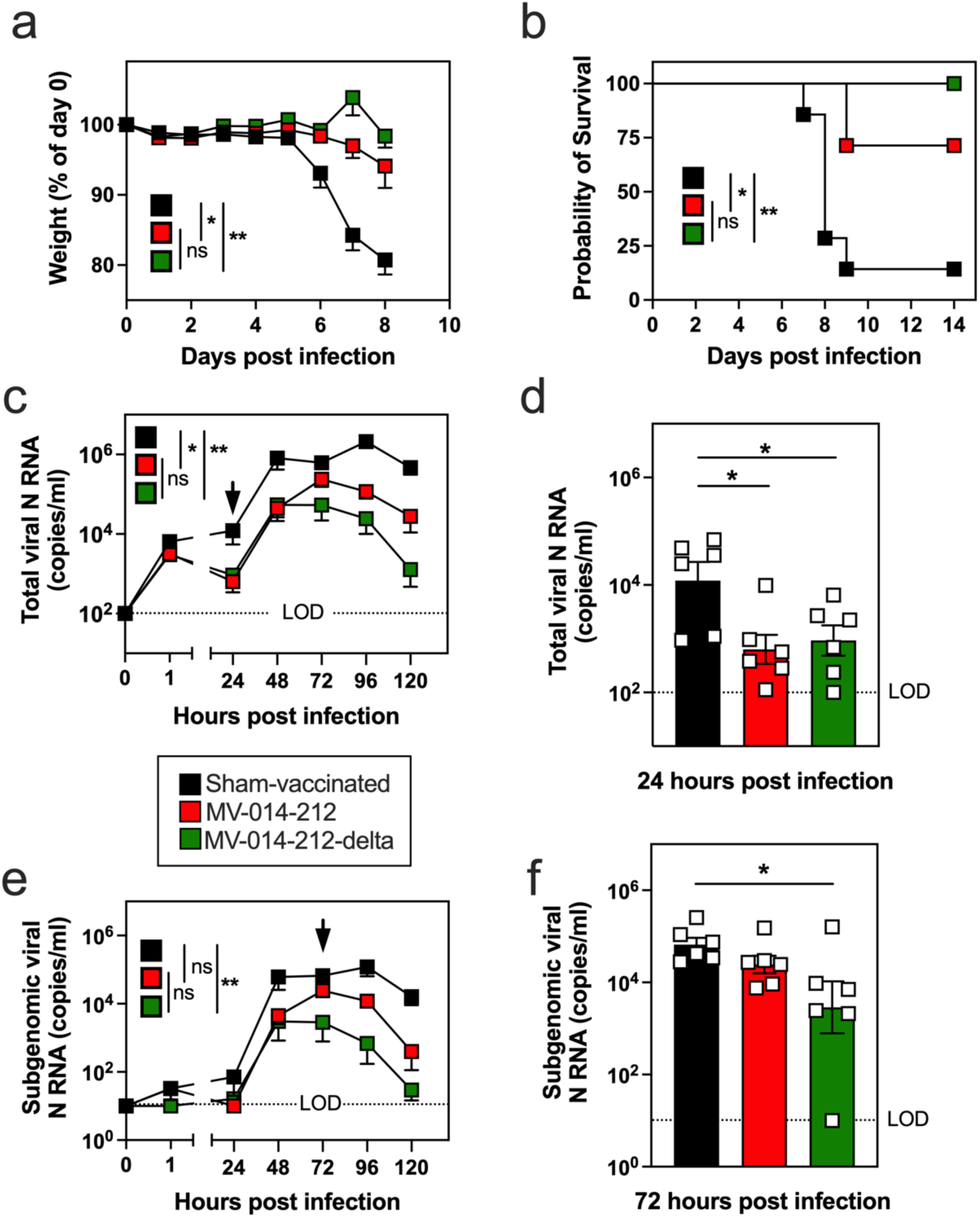
Morbidity, lethality and viral transcripts in nasal discharge from vaccinated mice following infection with Delta SARS-CoV-2. K18-hACE2 mice were sham-vaccinated (black) or vaccinated with MV-014-212 (red) or MV-014-212-delta (green) and infected with 2x10^4^ PFU Delta SARS-CoV-2 at day 35 post vaccination. Body weight loss measured as percent of initial weight (**a**) and survival (**b**) (n=7/group). Quantitation of total viral N RNA (**c**, **d**) and subgenomic viral N RNA (**e, f**) from nasal swipes collected longitudinally (daily) post infection. Panels **c** and **e** show longitudinal analysis from 0-120 hours post infection. Panels **d** and **f** show results from 24-hour (**d**) and 72-hour (**f**) timepoints post infection. Individual animals (n=6/group) plotted as symbols with group means shown as bars. All viral RNA data plotted as log-transformed copies/ml values. LOD, limit of detection. *P<0.05, **P<0.01. P values determined by one-way ANOVA followed by Dunnett’s multiple comparison test on area under the curve (AUC) measurements (**a**, **c**, and **e**), Log-rank (Mantel-Cox) test (**b**), or one-way ANOVA followed by Dunnett’s multiple comparison test (**d, f**). Variability expressed as SE.

Next, we measured viral shedding from the nasal passages of vaccinated/sham-vaccinated mice by quantifying total viral N RNA in nasal discharge isolated from nasal swipes collected daily following the challenge infection with Delta SARS-CoV-2. Total viral N RNA was detected as early as 1-hour post infection in the sham-vaccinated, MV-014-212 vaccinated and MV-014-212-delta vaccinated mice (**Figure 3c**), confirming successful i.n. instillation of the Delta SARS-CoV-2 challenge virus. Mice vaccinated with MV-014-212 or MV-014-212-delta had 1.4-1.5 log_10_ lower levels of total viral N RNA at 24 hours relative to the sham-vaccinated mice (**Figures 3c and 3d**). The vaccinated animals never reached the peak levels of viral N RNA seen in the sham-vaccinated mice and exhibited more rapid loss of the viral N RNA transcripts compared to sham-vaccinated mice (**Figure 3c**). As a surrogate measure of replicating SARS-CoV-2 virus, we next quantified subgenomic viral N RNA in the longitudinal collection of nasal swipes. Peak levels of subgenomic viral N RNA transcripts at 48 hours post-infection were 1.3-1.4 log_10_ lower in nasal swipes from vaccinated mice compared to sham-vaccinated mice (**Figure 3e**), and this decreased load of replicating virus transcripts was associated with more rapid reductions in subgenomic viral N copy number beginning at 72 hours post infection (**Figures 3e and 3f**). Consistent with the increased survival of the MV-014-212-delta vaccinated mice, we observed accelerated loss of both total and subgenomic viral N RNA transcripts in the mice immunized with the Delta Spike-expressing MV-014-212-delta vaccine compared to mice immunized with the USA/WA-1/2020 Spike-expressing MV-014-212 vaccine. These data show that both the MV-014-212 and MV-014-212-delta vaccines enhanced protection in mice infected with Delta SARS-CoV-2 and limited shedding of infectious virus from nasal secretions. However, while both vaccines provided protection from virus-induced morbidity and mortality, the vaccine that was antigenically matched to the challenge virus was, not surprisingly, the most protective.

### 2.4 A single i.n. administration of MV-014-212 or MV-014-212-delta vaccine attenuates pulmonary SARS-CoV-2 replication in mice

Intranasal infection of K18-hACE2 mice with SARS-CoV-2 results in infection that spreads from the nasal mucosa in the URT to the lungs in the LRT [43] and eventually to other organs, including the brain where the virus causes encephalitis and mortality [44]. To determine whether the RSV-vectored vaccines could protect from viral spread outside of the respiratory tract, we infected the sham-vaccinated and vaccinated mice with the SARS-CoV-2 Delta variant and then measured viral loads in both lung and brain tissues at day 5 post infection – a time when peak viral load in the lung is observed [43] and when virus can be detected in the brain [45]. As expected, mice that were sham-vaccinated and then infected presented with elevated levels of total viral N RNA transcripts in the lungs at day 5 post infection, ranging between 9-10 log_10_ copies/ml (**Figure 4a**). In contrast, total viral N RNA levels were significantly lower in the lungs of mice vaccinated with MV-014-212 or MV-014-212-delta compared to the sham-vaccinated mice. Importantly, subgenomic viral N RNA was not detected in the Day 5 post-infection lungs isolated from either group of vaccinated mice (**Figure 4b**). In contrast, 5/6 of the sham-vaccinated animals had detectable subgenomic viral N RNA in the lung (**Figure 4b**). Next, we measured infectious viral load in the lungs by plaque assay. All sham-vaccinated mice had viral titers above 4 log_10_ PFU/ml (**Figure 4c**). Conversely, both vaccinated groups of mice had significantly lower virus load in the lung with no measurable infectious virus present in 2/6 MV-014-212 vaccinated animals and 4/6 MV-014-212-delta vaccinated animals (**Figure 4c**).

**Figure 4.**
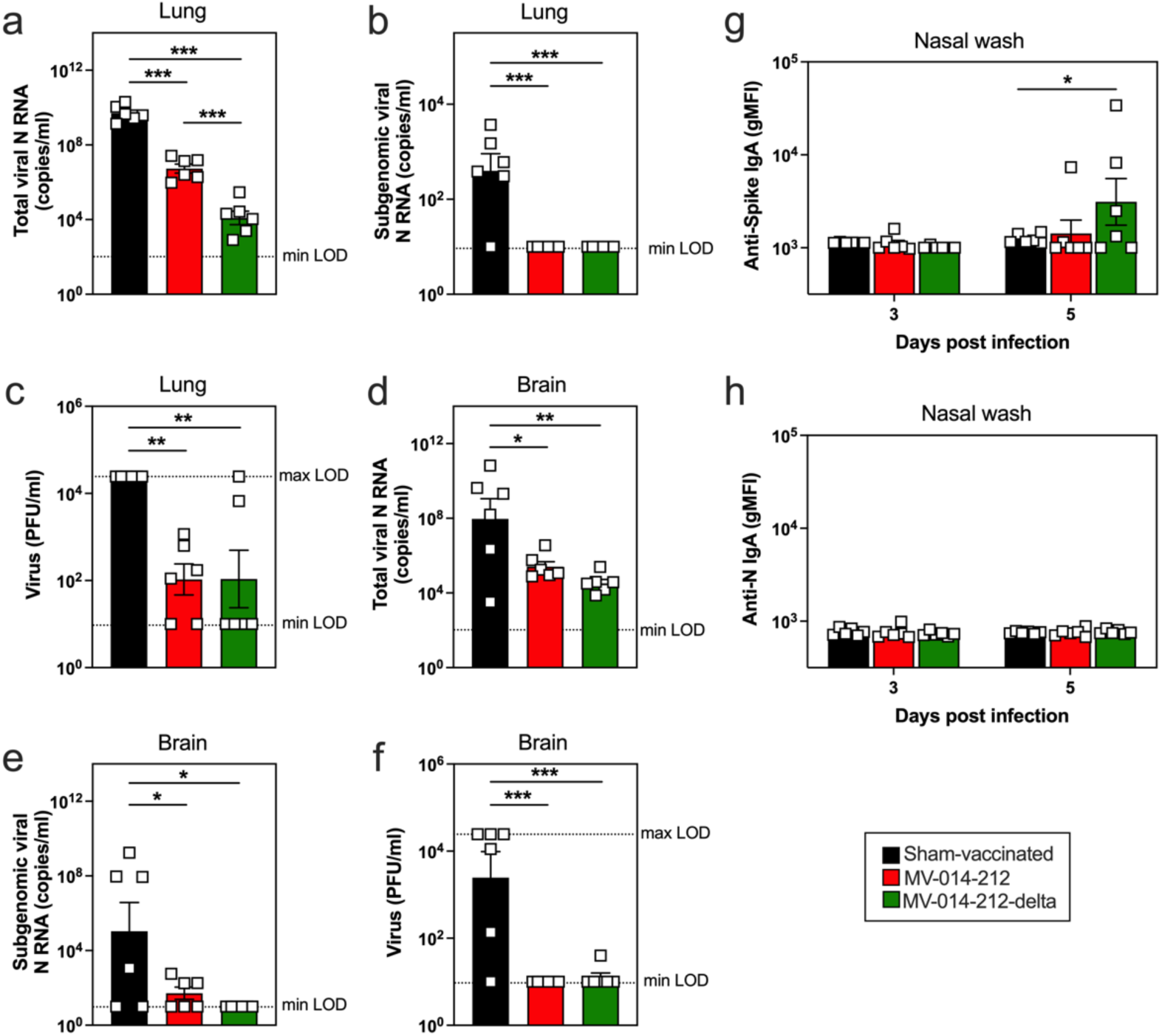
Viral loads in the lung and brain of vaccinated mice following infection with Delta SARS-CoV-2. Viral loads in lung (**a-c**) and brain (**d-f**) tissues from sham-vaccinated (black), MV-014-212 vaccinated (red), and MV-014-212-delta (green) vaccinated mice at day 5 post challenge infection with Delta SARS-CoV-2. Data shown as log-transformed copies/ml of total viral N RNA (**a, d**) and subgenomic viral N RNA (**b, e**), or as infectious virus measured by plaque assay and reported as log-transformed viral PFU/ml (**c, f**). Spike-specific (**g**) and N-specific (**h**) IgA antibodies in nasal wash from vaccinated K18-hACE2 mice at days 3 and 5 post infection with Delta SARS-CoV-2, reported as log-transformed geometric mean fluorescence intensity (gMFI). LOD, limit of detection. Individual animals (n=6/group) plotted as symbols with group means shown as bars. *P<0.05, **P<0.01, ***P<0.001. P values for **a-f** determined by one-way ANOVA followed by Dunnett’s multiple comparison test. P value for **g** determined by two-way ANOVA followed by Dunnett’s multiple comparison test. Variability expressed as SE.

To address whether the virus could be detected outside the respiratory tract, we assessed viral invasion of brain tissue. Consistent with previously published reports [45, 46], measurable virus was detected in the brain homogenate from sham-vaccinated mice with 4/6 brains from sham-vaccinated mice displaying high levels of total (**Figure 4d**) and subgenomic (**Figure 4e**) viral N RNA (≥8 log_10_ copies/ml) as well as infectious virus that was readily measured by plaque assay (**Figure 4f**). Conversely, mice vaccinated with MV-014-212 or MV-014-212-delta had measurable but limited levels of total viral N RNA in the brain (<6.5 log_10_ copies/ml) (**Figure 4d**), minimal to no subgenomic viral N RNA transcripts (**Figure 4e**) and no detectable infectious virus (**Figure 4f**). Therefore, i.n. vaccination with either MV-014-212 or MV-014-212-delta prevented spread of Delta SARS-CoV-2 to the brain.

In light of evidence that SARS-CoV-2 can travel to the brain from the nasal cavity via the olfactory bulb [47], we speculated that the efficacy of MV-014-212 and MV-014-212-delta vaccines at preventing SARS-CoV-2-induced encephalitis in mice likely stemmed from the vaccine-elicited induction of a protective nasal mucosa immune response. To test this, we collected nasal washes from sham-vaccinated mice and mice vaccinated with MV-014-212 or MV-014-212-delta at days 3 and 5 post-challenge with Delta SARS-CoV-2 and quantified Spike-specific IgA responses in the URT. No Spike-specific IgA was detected at day 3 post-infection in any group of mice (**Figure 4g**). However, by day 5 post-infection, half of the animals vaccinated with the Delta Spike-expressing MV-014-212-delta vaccine had detectable Spike-specific IgA in the nasal wash (**Figure 4g**). This modest Spike-specific IgA response appeared to be dependent on prior vaccination since Spike-specific IgA was not detected in nasal washes from the sham-vaccinated and then infected animals. Moreover, at this early timepoint, we did not detect IgA antibodies specific for viral antigens, like nucleocapsid (N), that were present in the infectious virus but not in the vaccine (**Figure 4g, 4h**). These data therefore argue that a single dose of the RSV-vectored vaccines, while not sufficient to induce a robust and durable IgA antibody-secreting cell (ASC) response in the URT, did appear to elicit memory B cells that could differentiate into mucosal IgA-producing ASCs following challenge infection with the same virus. In addition, the vaccine effectively decreased viral burden in the LRT and prevented virus spread to the brain.

### 2.5 A single i.n. administration of MV-014-212 or MV-014-212-delta vaccine protects hamsters against natural SARS-CoV-2 infection

Given the protective effects of MV-014-212 and MV-014-212-delta vaccines against Delta SARS-CoV-2 infection in K18-hACE2 mice and their effectiveness at inducing robust systemic and pulmonary immunity, we hypothesized that a single i.n. immunization with either vaccine would protect animals against naturally acquired infection mediated by co-housing vaccinated animals with recently infected animals. To test this, we moved to the Lakeview Golden (LVG) Syrian hamster model as these animals are highly susceptible to infection, able to support high levels of viral replication in the URT and LRT, and can generate fine aerosols that allow for transmission of respiratory viruses to cagemates [48–53].

To evaluate vaccine efficacy against naturally-acquired infection, hamsters were vaccinated i.n. with MV-014-212 or MV-014-212-delta, or were sham-vaccinated with Vero cell lysate. On day 28 post vaccination, a separate cohort of non-vaccinated, naive hamsters was experimentally infected with 2x10^6^ PFU Delta SARS-CoV-2. After 24 hours, individual experimentally-infected hamsters were paired with a single vaccinated or sham-vaccinated hamster. Following 24 hours of pairing, the cagemates were separated until termination of study (**Figure 5a**). To ensure that the hamsters were successfully immunized, we collected serum at day 24 and measured systemic Spike-specific IgG responses. As expected, the sham-vaccinated hamsters did not develop anti-Spike IgG (**Figure 5b**) or anti-RBD IgG (**Figure 5c**) antibodies in sera. Conversely, the hamsters vaccinated with MV-014-212 or MV-014-212-delta seroconverted and had increased levels of Spike and RBD-specific IgG by day 24 post vaccination (**Figures 5b and 5c**). Next, to assess morbidity in the experimentally infected hamsters and their vaccinated/sham-vaccinated cagemates, we measured body weights for 16 days following experimental infection (the infected hamsters) or following pairing (the vaccinated/sham-vaccinated cagemates). The experimentally infected hamsters began losing weight immediately following the i.n. infection with Delta SARS-CoV-2 and reached a maximum weight loss of 10-12% by day 6 post infection (**Figure 5d**). These animals did not recover their starting body weight, even 16 days later. Following pairing with experimentally-infected hamsters, the sham-vaccinated cagemates appeared to have become infected as evidenced by weight loss that peaked at 4-5% by day 6 post pairing (**Figure 5e**). The sham-vaccinated cagemates also failed to recover their starting body weight over 16 days (**Figure 5e**). For the first 4 days following pairing with experimentally-infected hamsters, the hamsters vaccinated with MV-014-212 or MV-014-212-delta lost weight. However, the weight loss did not surpass 2%, and the animals began regaining weight by day 5 post pairing and were fully recovered by day 13-14 post pairing (**Figure 5e**). Thus, a single i.n. dose of the RSV-vectored vaccine was not sufficient to prevent onset of morbidity in the vaccinated cagemates that were naturally infected by exposure to an infected hamster. However, vaccination strongly and significantly reduced morbidity and enabled a rapid and full recovery of body weight that was not observed in naturally infected, sham-vaccinated control animals over the same time-period.

**Figure 5.**
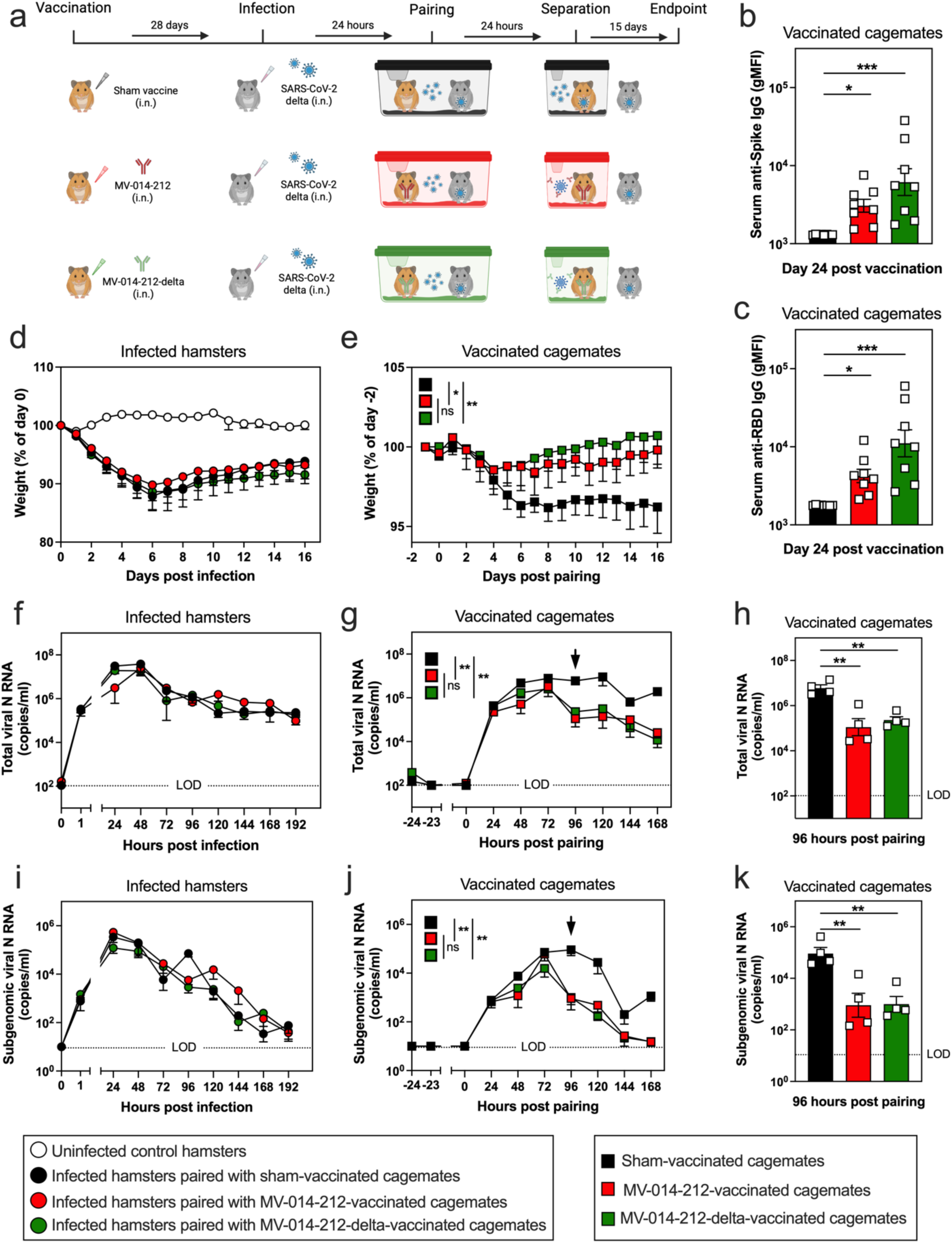
Morbidity and viral shedding from nasal discharge in vaccinated hamsters following naturally acquired SARS-COV-2 infection. (**a**) Experimental set-up for the SARS-CoV-2 naturally acquired infection study in LVG hamsters. Hamsters were vaccinated i.n. with Vero cell lysate (sham-vaccinated; black), MV-014-212 (3x10^5^ PFU; red) or MV-014-212-delta (3x10^5^ PFU; green). At day 28 post vaccination, a cohort of naïve hamsters was experimentally infected i.n. with 2x10^6^ PFU Delta SARS-CoV-2 (B.1.617.2). After 24 hours, the infected hamsters were paired (1:1) with a sham-vaccinated or vaccinated hamster for 24 hours to allow for natural transmission of Delta SARS-CoV-2 between the experimentally infected animal and its cagemate. The animals were then separated until study termination on day 16 post infection. (**b**) Spike-specific IgG and (**c**) RBD-specific IgG antibodies in sera at day 24 post vaccination with Vero cell lysate (sham-vaccinated; black), MV-014-212 (red) or MV-014-212-delta (green). Individual animals (n=8/group) shown as symbols and group means shown as bars. (**d, e**) Longitudinal body weight loss measurements of non-vaccinated hamsters that were experimentally infected with 2x10^6^ PFU Delta SARS-CoV-2 (**d**, circles) and in sham-vaccinated (black squares), MV-014-212 vaccinated (red squares) or MV-014-212-delta vaccinated (green squares) animals following pairing with the experimentally-infected hamsters (**e**). Body weight loss shown as percent of initial weight. Quantitation of total (**f-h**) and subgenomic (**i-k**) viral N RNA (n=4/group) in daily nasal swipes collected from experimentally-infected hamsters (**f, i**) and vaccinated/sham-vaccinated cagemates (**g, h, j, k**). Data reported as log-transformed copies/ml and shown for 0-192 hours post infection for experimentally-infected hamsters (**f, i**), for -24 to 168 hours post pairing for vaccinated/sham-vaccinated cagemates (**g, j**), or at 96 hours post pairing for vaccinated/sham-vaccinated cagemates (**h, k**). Data reported for individual animals (n=4/group) at the 96-hour time point, plotted as symbols with group means shown as bars. LOD, limit of detection. *P<0.05, **P<0.01, ***P<0.001. P values determined by one-way ANOVA followed by Dunnett’s multiple comparison test (**b-c**, **h**, **k**), and one-way ANOVA followed by Dunnett’s multiple comparison test on AUC measurements (**d-g**, **i, j**). Variability expressed as SE.

In addition to weight measurements, we also collected nasal swipes daily until day 8 (192 hours) post infection to quantify viral shedding in the nasal discharge. Total viral N RNA transcripts were detected in the experimentally-infected (non-vaccinated) hamsters as early as 1 hour post infection (**Figure 5f**), which confirmed a successful i.n. instillation of Delta SARS-CoV-2. Total viral load in the experimentally infected hamsters, as measured by total viral N RNA copies in nasal swipe samples, increased to 8 log_10_ copies/ml by 48 hours and then began to decline slowly, with 5-6 log_10_ copies/ml still detected at 192 hours post infection. Total viral N RNA was also detected within 24 hours of pairing the vaccinated/sham-vaccinated cagemates with experimentally infected hamsters (**Figure 5g**). No difference in total viral N RNA between vaccinated and sham-vaccinated animals was observed during the first 72 hours post pairing, but a significant decline in total viral N RNA transcript levels was observed starting at 96 hours post pairing in the hamsters vaccinated with MV-014-212 or MV-014-212-delta compared to the sham-vaccinated animals (**Figures 5g and 5h**).

Next, we quantified subgenomic viral N RNA in the nasal discharge to compare infectious viral load between the different groups of hamsters (**Figures 5i-k**). Subgenomic viral N RNA in the experimentally-infected (non-vaccinated) hamsters peaked at 5-6 log_10_ copies/ml at 24 hours post infection, began falling at 72 hours and declined to near limit of detection by 192 hours (**Figure 5j**). Subgenomic viral N RNA was also detected in the sham-vaccinated cagemates and the animals vaccinated with MV-014-212 or MV-014-212-delta following pairing with experimentally-infected hamsters (**Figure 5j**). For the first 72 hours following pairing, subgenomic N RNA levels in the nasal swab samples rose similarly between the sham-vaccinated and vaccinated cagemates (**Figure 5j**). However, the hamsters vaccinated with MV-014-212 or MV-014-212-delta showed a rapid drop in subgenomic viral N RNA at 96 hours post pairing that declined to background levels by 168 hours (**Figures 5j and 5k**). Conversely, the naturally infected, sham-vaccinated hamsters maintained maximal levels of subgenomic viral N transcripts until 120 hours post pairing with levels remaining elevated over background for at least 168 hours post pairing.

To better visualize the difference in viral clearance between the sham-vaccinated cagemates and the animals vaccinated with MV-014-212 or MV-014-212-delta, we compared the levels of total or subgenomic N RNA transcripts in nasal swipes from the experimentally infected hamsters to their vaccinated or sham-vaccinated cagemates. We first determined the number of total or subgenomic viral N RNA transcripts present in nasal swipes from the infected hamsters at day 5 post experimental infection and the number of total or subgenomic viral N RNA transcripts present in nasal swipes from the vaccinated/sham-vaccinated cagemates on day 5 post pairing. Next, we calculated the ratio of total N transcripts (**Figure S4a**) and subgenomic N transcripts (**Figure S4b**) for each pair of cagemates. As expected, the ratio of both total and subgenomic viral N RNA was significantly larger in the cages that co-housed experimentally-infected hamsters with sham-vaccinated cagemates when compared to cages co-housing experimentally-infected hamsters with vaccinated cagemates (**Figure S4a and S4b**).

Finally, to determine whether vaccination with MV-014-212 or MV-014-212-delta protected hamsters from lung injury due to naturally acquired infection with SARS-CoV-2, lung tissue sections were prepared from the sham-vaccinated and vaccinated hamsters on day 16 post pairing and then examined and scored by a board-certified pathologist. Lung histopathology was quantified by totaling the extent of damage to the alveoli (**Figure 6a-b**), airways (**Figure 6c-d**) and vasculature (**Figure 6e-f**). We found that 2/4 hamsters in both groups of vaccinated animals were fully protected from SARS-CoV-2-induced alveolar damage (**Figure 6a**). When the data from the two groups of vaccinated animals were combined to increase the size of the group, the alveolar damage score was significantly (P<0.05) reduced in the vaccinated hamsters compared to their sham-vaccinated counterparts (**Figure 6b**). Similarly, vaccinated hamsters had a decreased airway score compared to sham-vaccinated controls (**Figure 6c, 6d**). Conversely, no difference in vascular pathology was observed between vaccinated and sham-vaccinated hamsters, even when data from the two groups of vaccinated hamsters were combined (**Figure 6e, 6f**). Regardless, when all histology scores were taken in aggregate to determine a total histology score, the vaccinated hamsters had a significantly (P<0.05) lower total histology score compared to sham-vaccinated hamsters (**Figure 6g, 6h**). This lower score was primarily attributed to decreased prevalence of inflammation and compensatory epithelial changes affecting the alveolar airways, which were moderate (10-25% of alveolar tissue) in sham-vaccinated animals (**Figure 6i**) compared to minimal (less than 10% of alveolar tissue) in the vaccinated cohorts (**Figure 6j and 6k**). The inflammatory cells, which were predominantly mononuclear (lymphocyte, plasma cells, macrophages) infiltrated the alveolar interstitium, alveolar airways, and around vessels of sham-vaccinated hamsters (**Figure 6i**). Overall, these data suggest that the MV-014-212 and MV-014-212-delta vaccines attenuated virus-induced inflammation and injury to the alveolar and airway compartments of the lung.

**Figure 6.**
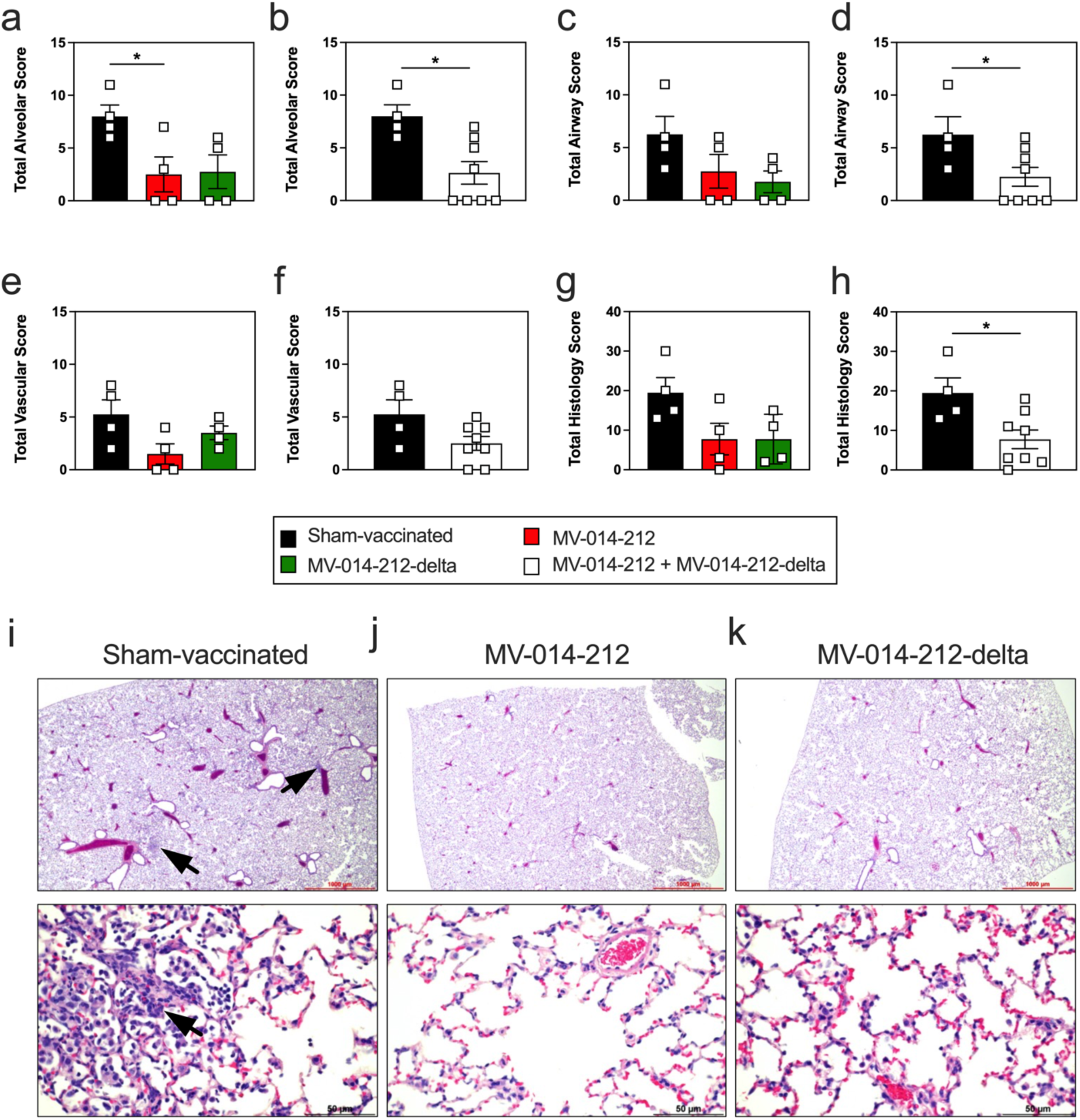
Lung inflammation and injury in vaccinated hamsters following natural SARS-CoV-2 infection. (**a-h**) Lung histology in sham-vaccinated hamsters (black) and hamsters vaccinated with MV-014-212 (red) or MV-014-212-delta (green) at day 16 post pairing with infected animals. Reported are alveolar histology scores (**a, b**), airway scores (**c, d**), vascular scores (**e, f**) and total histology scores (**g, h**). Comparisons include sham-vaccinated (black) to MV-014-212 (red) or MV-014-212-delta (green) vaccinated hamsters (**a, c, e, g**) or between sham-vaccinated hamsters (black) and all hamsters receiving a vaccine (white, panels **b, d, f, h**). (**i-k**) Representative H&E-stained images of lungs harvested from sham-vaccinated hamsters (**i**) and hamsters vaccinated with MV-014-212 (**j**) or MV-014-212-delta (**k**) at day 16 post pairing with infected animals. Images shown at 1,000 µm (top) and 50 µm (bottom) resolution. Inflammatory cell infiltrates are indicated by a black arrow. Individual animals (n=4-8/group) in **a-h** shown as symbols and group means shown as bars. *P<0.01. P values determined by one-way ANOVA followed by Dunnett’s multiple comparison test (**a, c, e, g**) or by two-tailed Student’s t-test (**b, d, f, h**). Variability expressed as SE.

Collectively, these results demonstrate that a single intranasal dose of RSV-vectored MV-014-212 or MV-014-212-delta vaccines elicited systemic and mucosal immunity and conferred protection against SARS-CoV-2 challenge, with MV-014-212-delta providing enhanced protection against the Delta variant. Furthermore, in naturally acquired infection studies, MV-014-212 and MV-014-212-delta reduced weight loss, limited viral replication within the nasal mucosa, and attenuated lung pathology in vaccinated hamsters exposed to infected cagemates, confirming the ability of these RSV-vectored vaccines to elicit broad protective respiratory immunity.

## 3. Discussion

We demonstrate that a single i.n. dose of a live-attenuated RSV-vectored vaccine encoding either the ancestral USA/WA-1/2020 Spike (MV-014-212) or the Delta Spike (MV-014-212-delta) elicits local and systemic immunity and significantly reduces SARS-CoV-2–induced morbidity and mortality following either experimental challenge infection or naturally acquired infection in preclinical animal models. These findings support the promise of intranasal mucosa-targeted vaccine strategies to generate front-line immunity at the portal of respiratory virus entry – an outcome that should reduce disease burden, decrease time to pathogen clearance following natural exposure to infected individuals, and lessen the risk of onward transmission by the vaccinated individuals who do become infected.

We observed a robust humoral response following vaccination that included systemic Spike-specific IgM antibodies that arose quickly and IgG antibodies that remained elevated for a longer duration. It is appreciated that early IgM responses can provide immediate, broad reactivity during the window before high-affinity neutralizing IgG antibodies develop and may contribute to early viral control at mucosal surfaces [54]. Indeed, several clinical studies report that delayed or absent early antibody responses, including delayed IgM seroconversion, are associated with more severe disease and worse COVID-19 outcomes [55–57]. Importantly, the systemic Spike-specific IgG antibodies elicited by the RSV-vectored vaccines, while arising later than the IgM antibodies, exhibited neutralizing activity. Interestingly, animals immunized with the Delta Spike (MV-014-212-delta) containing vaccine generated antibodies that neutralized virus expressing the ancestral USA/WA-1/2020 Spike protein, suggesting that the Delta Spike-containing vaccine elicited antibodies that could recognize shared or more broad neutralizing Spike determinants. Thus, a single dose of the RSV-vectored vaccine, even when administered via the mucosal route, appears sufficient to support layered humoral protection, with IgM acting rapidly and IgG mediating durable systemic neutralization and Fc-mediated effector functions.

Mucosal IgA is widely regarded as a critical correlate for neutralizing respiratory viruses at the site of entry and for blocking transmission [7]. While we expected that the mucosal delivery of the RSV-vectored vaccine would induce a strong IgA antibody response, we did not detect any Spike-specific IgA following vaccination. Instead, we detected modest Spike-specific IgA in nasal washes of MV-014-212-delta-vaccinated mice early following challenge infection. This response appeared to be vaccine-specific as no Spike-specific IgA at the early timepoints was observed in nasal wash from animals that were sham-vaccinated and then infected. In addition, the early IgA response, while elicited by the challenge infection, occurred before the *de novo* response to the challenge virus as antibodies to viral N protein, which was not present in the vaccine, were not observed at this early timepoint. Therefore, these data argue that the IgA Spike response observed early following challenge infection was almost certainly derived from the vaccine-elicited memory B cells. The fact that we only detected the mucosal IgA response in animals immunized with the delta Spike-containing vaccine and subsequently infected with Delta variant virus strongly suggests that the memory B cells needed to be activated by their cognate antigen in order to differentiate into IgA producing plasmablasts. We previously reported [12] that mucosal antigen delivery is required for the establishment of a lung mucosal resident memory B cell response. Likewise, mucosal antigen delivery is required for establishment of the URT IgA response [6, 18, 58, 59]. Therefore, the intranasal priming with the RSV-vectored vaccine likely initiated formation of the local lung resident memory B cell compartment that subsequently gave rise to local IgA producing antibody secreting cells (ASC) following antigen re-exposure. We would therefore argue that inclusion of an i.n. booster would maximize the nasal IgA-producing ASC and potentially achieve stronger immunity following exposure to live virus. Consistent with this, vaccination regimens that include i.n./i.n. prime+boost or systemic priming followed by i.n. boosters (prime/pull) amplify local IgA and resident immune populations and show promise in preclinical studies to enhance mucosal protection [35, 58, 60, 61].

Our data show that mucosal delivery of the RSV-vectored vaccine not only supported development of an URT IgA response following challenge but also elicited lung resident memory T cells that can contribute to rapid viral clearance following challenge infection. While these T cells may be more broadly reactive and capable of responding to multiple viral variants [62], T cell recall responses are not sterilizing and can reduce but not prevent infection [63, 64]. We found that the RSV-vectored vaccines expressing either the ancestral USA/WA-1/2020 Spike protein or the Delta variant Spike protein was sufficient to provide significant protection from morbidity and mortality following challenge infection. This result was consistent with published data [65–68] showing that vaccination does protect against severe disease even when the viral antigens are not matched between the vaccine and challenge virus. However, we also observed that when the vaccine Spike antigen was matched to the Spike expressed by the challenge virus, the protection following challenge infection was better. The most obvious explanation for the reduced protection in vaccinated animals challenged with heterologous virus was that the antibodies generated in these animals less effectively neutralized the heterosubtypic virus. Mutations in Delta (e.g., L452R, T478K) and similar mutations in other related SARS-CoV-2 strains are known to alter key neutralizing epitopes on Spike, which reduces binding by many vaccine-elicited antibodies and results in measurable reductions in neutralization titers [69–71]. These reductions in neutralization titers are commonly reported in the ∼2–6-fold range, depending on assay and serum source [72, 73]. However, even these relatively modest declines in neutralization can diminish vaccine effectiveness and result in increased symptom burden and disease severity. Together, the data argue that (i) antigenic matching or broader antigen designs are important for limiting early URT replication, and that (ii) mucosal vaccine strategies that generate local immunity (IgA and resident memory T cells) may mitigate the consequences of partial systemic neutralization loss by reducing infection at the portal of entry [59, 61, 74].

The real-world goal of vaccination is to prevent infection from occurring in the vaccinated individual after that individual is exposed to someone who is actively infected with the pathogen.

Reports describing testing other i.n. vaccine platforms in animals [61, 75], show decreased transmission between vaccinated and infected animals and naïve cagemates resulting in reduced onward spread after mucosal immunization. These experiments were performed as is typical for the laboratory setting where the vaccinated animal is given the challenge infection by direct administration of a single large bolus of live virus instilled in a large volume that results in simultaneous infection of both the URT and LRT of the animal. To examine whether the animals receiving the i.n. RSV-vectored vaccine were protected from a naturally acquired infection, we utilized LVG Syrian hamsters that are well known to support SARS-CoV-2 transmission through both direct contact and aerosols [76, 77]. We found that hamsters given a single i.n. vaccine with the RSV-vectored platform became infected when exposed to infected cagemates for a 24-hour period. Thus, the single dose of vaccine, which did not result in an ongoing local IgA response in the URT, was not sufficient to elicit sterilizing immunity and prevent productive infection as measured by levels of subgenomic viral N RNA transcripts in the URT over the first three days following a brief 24-hour period of cohousing with an infected cagemate. However, within 96 hours post pairing with an infected hamster, the vaccinated cagemates experienced reduced morbidity, accelerated viral clearance in the nose, and attenuated lung pathology compared to sham-vaccinated controls. Thus, the single administration of the vaccines, while not preventing naturally acquired infection, did significantly reduce the time in which productive virus could be found in nasal discharge. Again, it is possible that i.n. booster vaccination could further reduce the likelihood of the vaccinated cagemate becoming infected upon encounter with an infectious hamster.

Our findings should be interpreted within the context of the experimental framework used. The K18-hACE2 model provides a stringent and well-established system for evaluating severe disease and protection, though its ACE2 expression pattern differs from that of humans. Subsequently, extending these observations to additional models (e.g., non-human primates or alternative mouse systems) would further define translational applicability. We also observed vaccine-induced mucosal IgA in a subset of animals after challenge infection, albeit at modest levels, highlighting an opportunity to further optimize mucosal immunity through i.n. booster regimens, heterologous prime–pull approaches, or adjuvants optimized for enhanced mucosal induction. Finally, given the ongoing antigenic drift in SARS-CoV-2, next-generation designs incorporating conserved antigenic domains, multivalent or mosaic Spike constructs, or pan-sarbecovirus targets may further broaden the protective scope of the vaccine.

In conclusion, a single i.n. dose of MV-014-212 or MV-014-212-delta elicits coordinated systemic and mucosal immune responses that reduce disease burden and limit transmission in preclinical models. These findings underscore the potential of RSV-vectored i.n. respiratory pathogen vaccines as a practical strategy to couple durable systemic protection with front-line mucosal immunity. The results support continued optimization of i.n. boosting strategies with the vectors to strengthen mucosal IgA and tissue-resident memory responses, with the overarching goal of achieving sterilizing immunity and meaningful interruption of respiratory pathogen spread.

## 4. Materials and Methods

### 4.1 Design and generation of MV-014-212 and MV-014-212-delta vaccines

MV-014-212 and MV-014-212-delta are live-attenuated recombinant vaccines using a respiratory syncytial virus (RSV) backbone in which the native RSV G and F surface glycoproteins were replaced with either a chimeric SARS-CoV-2 USA-WA1/2020 Spike protein (MV-014-212) or a chimeric Delta SARS-CoV-2 (B.1.617.2) Spike protein (MV-014-212-delta) comprising the Spike ectodomain and transmembrane domain fused to the cytoplasmic tail of RSV F (strain line 19). The design, attenuation strategy, and preclinical characterization of this RSV vaccine platform been described previously [38].

### 4.2 Animals

K18-hACE2 transgenic mice [39] that express human angiotensin-converting enzyme 2 (hACE2), the primary host cell receptor for severe acute respiratory syndrome (SARS) coronavirus [78], were purchased from The Jackson Laboratory (strain # 034860) and bred at The University of Alabama at Birmingham (UAB). Lakeview Golden (LVG) Syrian hamsters were purchased from Charles River Laboratories.

### 4.3 Vaccinations

8-12-week-old K18-hACE2 mice and 12-week-old male LVG Syrian hamsters were vaccinated intranasally with the live-attenuated, RSV-vectored vaccines described above. Animals were sedated with isoflurane (3-4% with 0.3-0.5L/minute (min) O_2_) and inoculated while in a supine position. Mice were intranasally inoculated with 1.5x10^5^ PFU vaccine in a 50 μl volume. Hamsters were intranasally inoculated with 3x10^5^ PFU vaccine in 100 μl volume (50 μl/nare). Sham-vaccinated mice and hamsters were intranasally inoculated with 50 μl and 100 μl, respectively, of Vero cell lysate.

### 4.4 Serum collections

Blood samples from mice were collected via submandibular vein puncture in awake animals and via the saphenous vein in sedated hamsters. Terminal blood samples were obtained from both species by vena cava collection. All samples were stored into BD Microtainer® tubes (BD Biosciences) and centrifuged at 10,000 × *g* for 10 min at room temperature (RT). Sera were separated and, when collected post-infection, heat-inactivated at 60 °C for 20 min before removal from the ABSL-3 containment facility. Serum was aliquoted and stored at -80°C until analyzed.

### 4.5 Production of recombinant SARS-CoV-2 proteins

Recombinant SARS-CoV-2 Spike ectodomain trimers were generated from two human codon-optimized constructs with the following linear sequence: a human IgG leader peptide, the Spike ectodomain (amino acids 14–1211) from the SARS-CoV-2 Wuhan-Hu-1 strain, a GGSG linker, the T4 fibritin foldon trimerization domain, a GS linker, and either an AviTag (construct 1) or a 6×His tag (construct 2). Both constructs incorporated two sets of stabilizing mutations to maintain the pre-fusion conformation: (i) substitution of the furin cleavage site RRAR→SGAG (residues 682–685) [79] and (ii) two proline substitutions (K983P, V984P) [79, 80]. To generate recombinant Spike receptor-binding domain (RBD), a separate human codon-optimized construct encoding a human IgG leader peptide, Spike RBD (amino acids 319–541 from Wuhan-Hu-1), a glycine linker, AviTag, and 6×His tag was used; no mutations were introduced in this construct.

Recombinant Spike ectodomain protein trimers were expressed in *FreeStyle™ 293-F* cells. Spike trimers were generated by co-transfection of Avi- and His-tagged plasmids at a 1:2 ratio. Monomeric RBD proteins were generated by transfection of the expression plasmid into *FreeStyle™ 293-F* cells. Transfected cultures were incubated for three days, and clarified supernatants were collected by centrifugation. Proteins were purified from the supernatant using fast protein liquid chromatography (FPLC) on a HisTrap HP column (GE Healthcare) with elution in 250 mM imidazole. Purified Spike trimers and RBD monomers were buffer-exchanged into either 10 mM Tris-HCl (pH 8.0) or 50 mM Bicine (pH 8.3) and subsequently biotinylated using biotin-protein ligase (Cytiva).

Recombinant coronavirus nucleocapsid (N) protein was produced from a human codon-optimized construct encoding a human IgG leader sequence, full-length N (Gene Bank accession #QHD43423), an AviTag, and a 6×His tag. The Avi-His–tagged N protein was expressed in *E. coli* Rosetta cells that were co-transfected with the N expression plasmid and an inducible *BirA* ligase plasmid. Cultures were grown in the presence of chloramphenicol, ampicillin, and streptomycin, followed by induction with biotin. Biotinylated N protein was purified using nickel-affinity FPLC and subsequent size-exclusion chromatography.

### 4.6 Generation of SARS-CoV-2 Spike B cell tetramers

Biotinylated full-length SARS-CoV-2 Spike protein (USA/WA-1 strain) was tetramerized using fluorochrome-conjugated streptavidin (PE or APC; Agilent). The optimal protein:streptavidin ratio required to achieve near-complete occupancy of streptavidin biotin-binding sites was first empirically determined based on estimated molar ratios and confirmed by Western blot titration. For tetramer assembly, streptavidin-fluorochrome was added stepwise to the biotinylated Spike protein (in ∼10% increments every 10 min) at room temperature with protection from light to minimize aggregation and promote uniform complex formation. Following conjugation, tetramers were diluted in 0.22 μm–filtered PBS to the desired working concentration, clarified by centrifugation at 20,000 × g for 10 min at 4°C, and the supernatant was collected. Sodium azide was added to a final concentration of 0.05%, and tetramers were aliquoted and stored at 4°C until use.

### 4.7 Generation of SARS-CoV-2 cytometric bead array

To generate the cytometric bead array (CBA), 500 µg of biotinylated antigen was conjugated to 2x10^7^ Streptavidin (SA)-functionalized 4 µm Carboxy Blue fluorescent micro beads (Spherotech CPAK-4067-8K) as previously described [81].

### 4.8 Measurement of SARS-CoV-2-specific antibody responses by cytometric bead array

Mouse and hamster sera (diluted 1:4000) and undiluted mouse nasal wash samples in 40 µL PBS were arrayed in 96-well U-bottom polystyrene plates. Each sample was combined with a 5-µL bead suspension containing 5 × 10⁵ SARS-CoV-2 Spike-conjugated beads, 5 × 10⁵ RBD-conjugated beads, 5 × 10⁵ coronavirus N-conjugated beads, and either 5 × 10⁵ beads conjugated to mouse anti-IgG, anti-IgM, and anti-IgA antibodies (mouse array) or 5 × 10⁵ beads conjugated to hamster anti-IgG antibodies (hamster array). Suspensions were mixed by gentle pipetting and incubated for 15 min at RT. Following incubation, beads were washed with 200 µL PBS and pelleted by centrifugation at 2100 × *g* for 10 min at RT. Bead pellets were resuspended in a secondary staining solution containing polyclonal goat anti-mouse IgG-Alexa Fluor 488 (1:400), anti-mouse IgM-PE (1:400), or anti-mouse IgA-PE (1:400; SouthernBiotech), or polyclonal anti-hamster IgG-FITC (1:400; SouthernBiotech). Samples were incubated for 15 min in the dark at RT, washed with 200 µL PBS, and centrifuged at 2100 × *g* for 10 min. Beads were resuspended in 100 µL 1% paraformaldehyde (PFA) and analyzed on a *CytoFLEX* flow cytometer (Beckman Coulter) operated in plate mode at a sample rate of 50 µL min⁻¹. Acquisition was stopped for each sample following 100 µL collection. Flow cytometry standard (FCS) files were analyzed using *FlowJo* software V10.10.0 (FlowJo LLC). Beads were identified by gating on singlet 4-µm particles in logarithmic forward- and side-scatter plots. Distinct bead populations were separated using APC-A750 fluorescence gates, and geometric mean fluorescence intensity (gMFI) was quantified in the PE and AF488/FITC channels.

### 4.9 Propagation and titer determination of Delta SARS-CoV-2

The SARS-CoV-2 Delta variant (B.1.617 clade) clinical isolate was obtained from a patient sample collected at the UAB Hospital. Virus was propagated in *Vero E6* cells (ATCC C1008) and titrated by viral plaque assay as previously described [46]. All procedures involving infectious SARS-CoV-2 were conducted in the Southeastern Biosafety Laboratory (SEBLAB) Biosafety Level 3 (BSL-3) facility at UAB in accordance with institutional biosafety protocols.

### 4.10 Infections with Delta SARS-CoV-2

LVG Syrian hamsters were infected i.n. with 2×10^6^ plaque-forming units (PFU) of the SARS-CoV-2 Delta variant in 100 µL phosphate-buffered saline (PBS; 50 µL/nare). Mice were infected i.n. with 2×10^4^ PFU of SARS-CoV-2 Delta variant in 30 µL PBS.

### 4.11 Nasal swipe collections

The exterior surface of each mouse and hamster nose was swabbed for 5–10 seconds (s) using a polyester-tipped applicator (Medical Packaging SP-7D) pre-moistened in 300 µL of viral transport medium (VTM; Hanks’ balanced salt solution [HBSS] containing Ca²⁺ and Mg²⁺, supplemented with 2% fetal bovine serum [FBS], 100 µg/mL gentamicin, and 0.5 µg/mL amphotericin B). Following sampling, the swab tip was aseptically cut and placed into the original VTM tube. After collection of all samples, tubes were vortexed briefly to release material, and swab tips were discarded. Nasal swipe samples were stored at -80°C until analyzed.

### 4.12 Nasal wash collections

Nasal washes were collected by flushing the nasal passages with 400 µL of VTM using a 19-gauge blunt needle attached to a 1-mL insulin syringe. Briefly, the trachea was incised following euthanasia, and the blunt needle was gently inserted. While the mouth was held closed, VTM was slowly flushed through the nasal passages, exiting through the nares into a sterile collection tube. Nasal wash samples were stored at -80°C until analyzed.

### 4.13 Tissue collection and processing for viral RNA quantitation

Mouse lungs and brains were excised postmortem and placed into 2-mL Lysing Matrix M tubes (MP Biomedicals, #116923050-CF) containing 1 mL VTM. Tissues were homogenized using a FastPrep-24 Classic bead-beating system (MP Biomedicals) at 4.0 m/s for 20 s. Homogenates were placed on ice for 5 min and homogenized a second time under identical conditions. The resulting lysates were centrifuged at 1000 x *g* for 10 min at RT, and clarified supernatants were collected and stored at -80°C until analyzed.

### 4.14 Tissue collection and processing for flow cytometry

Mouse lung tissue was harvested postmortem, mechanically minced, and digested in RPMI-1640 medium containing 1.25 mg/mL collagenase (MilliporeSigma) and 150 U/mL DNase I (MilliporeSigma) for 30 min at 37 °C. Digested tissue was passed through a 70-µm Falcon™ cell strainer (Fisher Scientific), pelleted by centrifugation, and resuspended in 3 mL red blood cell lysis buffer (10 mM KHCO₃, 150 mM NH₄Cl, 0.1 mM EDTA; pH 7.2–7.4). After incubation for 5 min at RT, lysates were diluted 1:5 in staining medium (SME; Dulbecco’s PBS supplemented with 2% FBS and 2 mM EDTA). Samples were filtered through a 100-µm Nitex® nylon mesh (Fisher Scientific), washed, and resuspended in SME. Mediastinal lymph nodes (mLNs) were harvested postmortem, gently dissociated through a 70-µm Falcon™ cell strainer, pelleted by centrifugation, and resuspended in SME.

### 4.15 Flow cytometry

Cell concentrations from each single-cell suspension were determined by adding 20 µL of sample into a 96-well V-bottom plate containing 50 µL of Fluoresbrite® carboxylate YG 10-µm microspheres (2.0 × 10⁵ beads/mL; Polysciences Inc.) and 180 µL of 7-AAD (1:720 dilution) prepared in SME. Samples were analyzed using a FACSCanto II flow cytometer (BD Biosciences) to quantify live cell numbers.

For immunophenotyping, 200 µL of each single-cell suspension was transferred to a 96-well V-bottom plate and incubated with Fc Block (10 µg/mL; clone 24G2) for 10 min at 4 °C. Following 200 µL SME wash, cells were stained in the dark at 4 °C for 20 min with either the B cell or CD8⁺ T cell phenotyping panel.

The B cell phenotyping panel included: CD19-APC-Cy7 (Biolegend, #115558, 1:200), CD138-BV421 (Biolegend, #142508, 1:200), IgD-BV510 (Biolegend, #405723, 1:500), CD38-PE-Cy7 (eBioscience, #25-0381-82, 1:400), CD95/FAS-FITC (BD, #554257, 1:200), F4/80-PerCP-Cy5.5 (Biolegend, #123130, 1:200), CD3e-PerCP-Cy5.5 (Invitrogen, #45-0031-82, 1:200), 7-AAD (Biolegend, #139308, 1:1000), SARS-CoV-2 Spike-PE tetramer (1:50) and SARS-CoV-2 Spike-APC tetramer (1:50). Samples were acquired on a FACSCanto II flow cytometer (BD Biosciences).

The CD8⁺ T cell phenotyping panel included: CD69-FITC (BD, #553236, 1:200), CD8α-BV605 (Biolegend, #100744, 1:200), CD45.2-BUV395 (BD Horizon, #564616, 1:200), CD4-BUV563 (BD, #612923, 1:200), CD3-BUV661 (BD, #750638, 1:200), CD103-BV711 (BD Horizon, #564320, 1:200), Aqua Live/Dead (ThermoFisher Scientific, #L34965, 1:1000), and Class I MHC H-2Kᵇ SARS-CoV-2 Spike (VNFNFNGL)-BV421 tetramer (NIH Tetramer Core Facility, Atlanta, GA; #63894, 1:100) Samples were acquired on a FACSymphony flow cytometer (BD Biosciences) within the UAB Flow Cytometry and Single Cell Core Facility.

All flow cytometry data were analyzed using FlowJo software (FlowJo LLC, Version 10.10.0 or later).

### 4.16 Viral RNA quantitation by qRT-PCR

Total and subgenomic SARS-CoV-2 N RNA levels were quantified by qRT-PCR as previously described [45]. Each homogenized sample was diluted 1:1 with lysis buffer containing 10% proteinase K from the Maxwell® RSC Viral Total Nucleic Acid Purification Kit (Promega), vortexed, and incubated at 56 °C for 10 min. RNA was extracted using the Maxwell® RSC 48 Instrument (Promega) according to manufacturer’s instructions. Quantitative RT-PCR was performed on a QuantStudio™ 5 Real-Time PCR System (ThermoFisher Scientific).

For genomic N RNA quantification, primers targeting the N gene (forward, 5′-GAC CCC AAA ATC AGC GAA AT-3′; reverse, 5′-TCT GGT TAC TAC TGC CAG TTG AAT CTG-3′) and a fluorogenic probe (/SFAM/ACC CCG CAT TAC GTT TGG TGG ACC/BHQ_1) were used (Integrated DNA Technologies, IDT). Viral RNA copy number was determined using a standard curve generated from AccuPlex™ SARS-CoV-2 Reference Material (SeraCare Life Sciences, #0505-0126) extracted and amplified in parallel.

For subgenomic N RNA quantification, a forward primer targeting the leader sequence (5′-CGA TCT CTT GTA GAT CTG TTC TC-3′; IDT) and the same N gene reverse primer were used. Amplification was performed using the Power SYBR™ Green RNA-to-CT™ 1-Step Kit (ThermoFisher Scientific). The sgRNA standard was constructed in-house by amplifying the N gene with primers sgRNA-UF (5′-CCA ACC AAC TTT CGA TCT CTT GTA-3′) and sgRNA-NR1 (5′-AAG GTC TTC CTT GCC ATG TTG-3′) (IDT). The amplified product was purified from agarose gel and quantified (2 × 10¹³ copies/mL) using a NanoDrop™ One Microvolume UV-Vis Spectrophotometer (ThermoFisher Scientific). Serial dilutions of this standard were used to generate a calibration curve for sgRNA quantitation.

### 4.17 Production of SARS-CoV-2 pseudovirus VSV-ΔG-Spike

Similar to published protocols [82], Vesicular stomatitis virus (VSV)-ΔG-G pseudovirus was generated at UAB using the G-VSV pseudotyping expression system, which utilizes the recombinant genomic plasmid, (rVSV)-ΔG-GFP-2.6 (Kerafast, Cat. EH1027), T7-based support plasmids (pVSVN, pVSVP, pVSVG and pVSVL; Kerafast, Cat. EH1012) and pCAGGS-VSVG plasmid (Kerafast, Cat. EH1017). Briefly, BSR-T7 cells were transfected with genomic and T7 support plasmids (N, P, G, L; ratio 3:5:8:1) in TransIT-LT1 (Mirus Bio, Cat. MIR 2300). The initial VSV-ΔG-G pseudotyped-virus was harvested from media. To amplify the virus, HEK293T cells were transfected with the pCAGGS-VSVG plasmid. 24 hours later, cells were infected with VSV-ΔG-G pseudotyped-virus at an MOI up to 0.1. Virus was harvested after 24 hours from media by centrifugation. To generate VSV-ΔG-Spike pseudovirus, adherent HEK293T cells were transfected with of the SARS-CoV-2 (2019-nCoV) Spike plasmid, where the C-terminal 21 amino acids of Spike were removed [83]. After 24 hours, cells were infected with VSV-ΔG-G pseudotyped-virus (MOI: 1.0). Supernatants containing the produced VSV-ΔG-Spike pseudovirus were harvested 24 hours post transfection and clarified by low-speed centrifugation. Supernatants which contain the concentrated pseudovirus were aliquoted and stored at -80 °C until used.

### 4.18 Pseudoneutralization assay

Twenty-four hours before performing the assay, 2.5 × 10⁴ human ACE2 (hACE2)-expressing HEK293T cells in 100 µL of complete medium were seeded per well in a 96-well flat-bottom culture plate. The following day, three-fold serial dilutions of serum samples (67 µL/well, in duplicate) were prepared in DMEM high-glucose medium supplemented with 5% FBS. Each dilution was mixed with an equal volume of pseudovirus-containing medium (2000 focus-forming units [FFU]/well of rVSV-ΔG-Spike). The virus–serum mixtures were incubated in the dark for 30 min at RT before being added (100 µL/well) to the plate containing adherent hACE2-HEK293T cells. Plates were incubated for 18–24 hours at 37 °C in 5% CO₂. Following incubation, cells were detached with trypsin, washed, and resuspended in SME buffer. GFP fluorescence, indicative of infection, was quantified on a CytoFLEX flow cytometer (Beckman Coulter). Data normalization was calculated by setting control wells containing virus and cells without sera (PBS only) as 0% neutralization.

### 4.19 Plaque assay

Vero E6 cells were seeded in 6-well plates at a density of 4 × 10⁵ cells per well and incubated for 16 hours at 37 °C in 5% CO₂ to allow monolayer formation. Serial tenfold dilutions of clarified lung homogenate supernatants were prepared in 1× PBS containing 1% FBS and added to the cell cultures. Plates were incubated for 1 hour at 37 °C with gentle agitation every 10 min to facilitate viral adsorption. Following incubation, inoculum was removed, and cells were overlaid with a medium containing 0.6% Avicel, 3% FBS, and 1× MEM. The plates were returned to the CO₂ incubator and maintained for 72 hours, afterwhich the Avicel overlay was carefully aspirated, and cell monolayers were fixed with 10% neutral-buffered formalin (NBF) for 1 hour at RT. Fixed cells were subsequently stained with 0.05% Neutral Red dye, and visible plaques were enumerated for each dilution. Viral titers were calculated by multiplying the number of plaques by the corresponding dilution factor. The lower limit of quantification for this assay corresponded to the lowest dilution of the plated lung homogenate.

### 4.20 Histology

Hamster lungs were inflated with 3 ml 10% NBF using a blunt 19-gauge needle attached to a 5-ml syringe. The tissues were collected and fixed for seven days in 50-ml conical tubes filled with 10% NBF. The fixed tissues were embedded dorsal side down in paraffin. Tissue blocks were sectioned at 5 µm, and slides containing the tissue sections were stained with hematoxylin and eosin (H&E). Tissue slides were evaluated by light microscopy by a board-certified veterinary pathologist blinded to study groups who assessed lung histopathologic lesions using a published scoring matrix [84]. Alveolar pathology in affected areas was scored using standardized reporting criteria [85]. Briefly, a total alveolar score (scale 0-4) was generated by individually assessing and summing the extent of alveolar inflammation, alveolar damage and type II pneumocyte hyperplasia. Representative images were taken using a Nikon Eclipse Ci microscope (Nikon Inc) and analyzed with NIS-Elements software (Nikon Inc). All Images were saved as JPGs and exported into GraphPad Prism (v10.3.0) for presentation. No post-acquisition image processing or adjustments (including contrast, brightness, or color) were performed.

### 4.21 Quantification and statistical analysis

Statistical methods, sample sizes (n), and the number of experimental replicates for each experiment are detailed in the corresponding figure legends. Statistical analyses and data visualization were performed with GraphPad Prism (v10.3.0).

### 4.22 Ethics statement

All procedures involving animals were reviewed and approved by the UAB Institutional Animal Care and Use Committee (IACUC), under protocol number 22559 that was approved on 1 June 2022.

## Acknowledgements

The authors would like to thank Uma Mudunuru, Thomas ‘Scott’ Simpler, Rebecca Burnham, Sara Ellen Callahan and Kelsey Browning for animal husbandry in ABSL2; Donovan J. Murphy and Robert Alldredge for animal husbandry in ABSL3; Moustafa A. Awaden, Justin C. Roth, Bryan LaGory and Vineel P. Reddy for support in the UAB SEBLAB; Derek B. Moates and the UAB Fungal Reference Lab for viral RNA quantitation; Emily E. Helmen and Sherri Coffman for histology support, and Andrea J. Osborne, DVM, MS, DACLAM for veterinary care of animals. Meissa Vaccines Inc. provided funding for the work reported in this article. Support for the development and validation of SARS-CoV-2 cytometric bead arrays was provided by U19 AI142737 and U19 142737-04S1 (FEL, RGK, TJG, TDR). Support for the UAB Flow Cytometry and Single Cell Core was provided by NIH P30 CA013148 and P30 AI027767. SEBLAB funding was provided by NIAID Regional Biocontainment Lab (RBL) awards UC6AI058599, 1G20AI167409-01, 1G20AI167409-S1 and 1UC7AI180255-01. The Emory University NIH Tetramer Core Facility (RRID:SCR_026557), which provided the Class I MHC H-2Kᵇ SARS-CoV-2 Spike (VNFNFNGL)-BV421 tetramer used in this study, is supported by NIH Contract 75N93020D00005. We also thank the UAB Immunology Institute Antibody Characterization and Serology Core (<u>UAB II ACS</u>) for providing the recombinant SARS-CoV-2 Spike and RBD proteins and B cell tetramers used in this study.

## Author Contributions

FEL, RST and MLM conceived the idea for the project and secured funding. FEL, RST, MLM, and TDR were responsible for supervision of personnel associated with the project. FEL, RST, MLM, DB and MDS designed the experiments that were performed by DB, MDS, ASS, and DK. MFT, CMW, and SIP contributed to RSV virus design, rescue, growth, harvest optimization and/or *in vitro* characterization. GY and JA generated the plasmids and expressed the SARS-CoV-2 antigens. RGK developed the CBA assay that was performed by JA and FZ. KSH provided the Delta SARS-CoV-2 virus. JLT performed the plaque assays with assistance from CAB. TJG and QS produced and provided the SARS-CoV-2 pseudovirus VSV-ΔG-Spike. LS and SML performed the viral RNA quantitation and analyzed data. JBF performed the histology analyses. All other data were analyzed by DB, MDS and FEL. DB, MDS and FEL wrote the manuscript and prepared the final figures. All authors read and approved the final manuscript.

