## Supplemental Figures for "Intranasal immunization with live-attenuated RSV-vectored SARS-CoV-2 vaccines elicits antigen-specific systemic and mucosal immunity and protects against viral challenge and natural infection"

**Supplemental Figures and Legends**

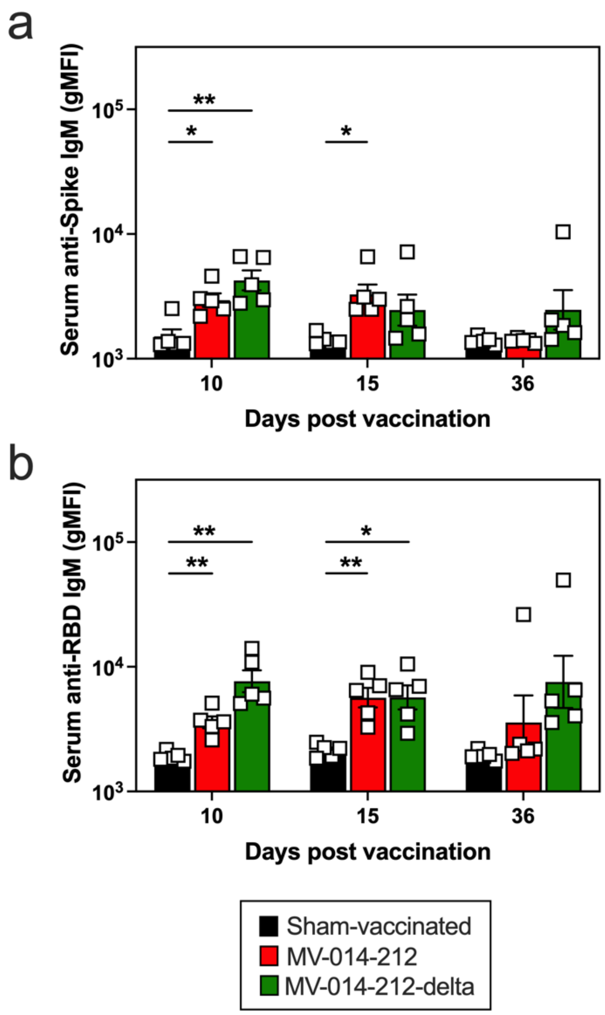


**Figure S1. Systemic antigen-specific IgM responses in mice following a single i.n. administration of MV-014-212 or MV-014-212-delta vaccine.** Spike-specific IgM (**a**) and RBD-specific IgM (**b**) in sera from K18-hACE2 mice at days 10, 15 and 36 post vaccination with Vero cell lysate (sham vaccine; black), MV-014-212 (red) or MV-014-212-delta (green). Data plotted as log-transformed geometric mean fluorescence intensity (gMFI). Individual animals (n=5/group) shown as symbols and means of groups shown as bars. *P <0.05, **P<0.01. P values determined by one-way ANOVA followed by Dunnett’s multiple comparison test. Variability expressed as standard error (SE).

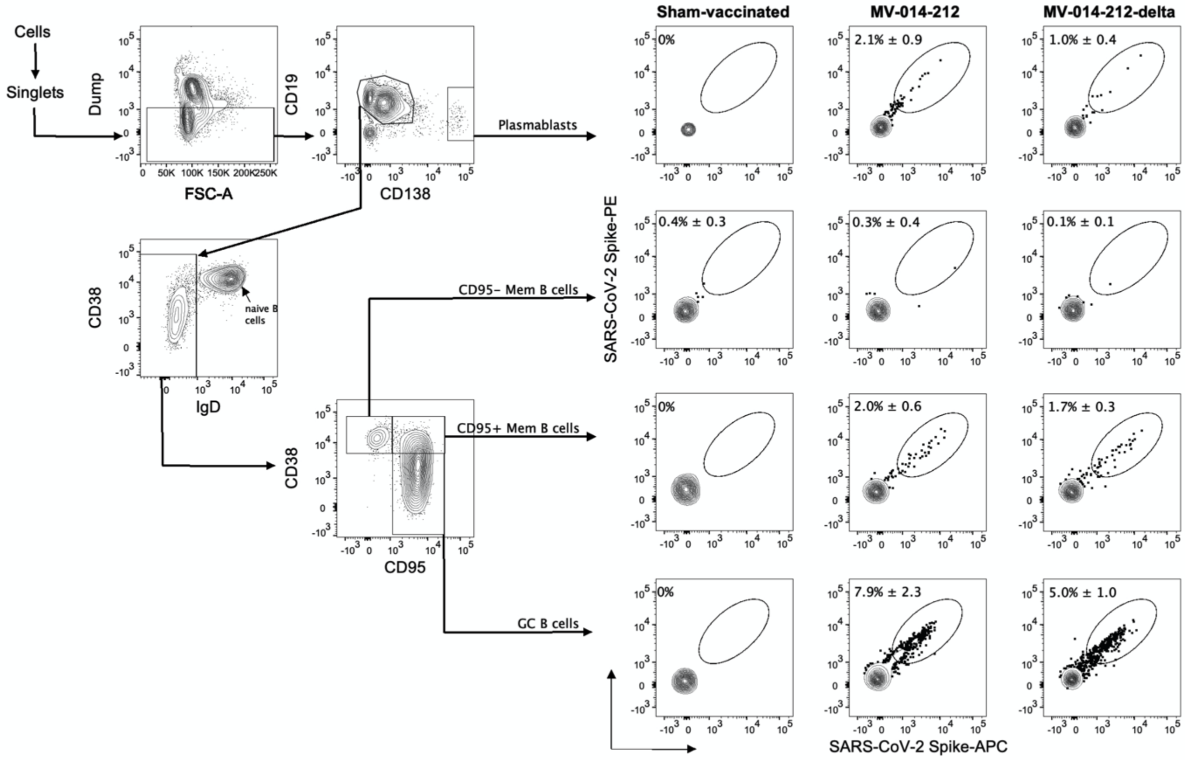


**Figure S2. Gating strategy for flow cytometric analysis of Spike-specific B cells in the mediastinal lymph node from vaccinated mice.** Sequential gating strategy for identification and enumeration of Spike-specific plasmablasts, CD95$^{+}$ memory (Mem) B cells, CD95$^{neg}$ Mem B cells and germinal center (GC) B cells in the mediastinal lymph node of K18-hACE2 mice at day 10 post vaccination with Vero cell lysate (sham-vaccinated), MV-014-212 or MV-014-212-delta. A dump channel containing 7-Aminoactinomycin D (7-AAD) viability dye, anti-mouse F4/80 and anti-mouse CD3 antibodies was used to exclude dead cells, macrophages and T cell lymphocytes from the analysis. Plasmablasts were phenotyped as Dump$^{neg}$ CD19$^{+/lo}$ CD138$^{+}$ cells. Mem B cells were phenotyped as Dump$^{neg}$ CD19$^{+}$ IgD$^{neg}$ CD38$^{hi}$ cells. Mem B cells were further subdivided into CD95$^{neg}$ and CD95$^{+}$ Mem B cell subsets. GC B cells were phenotyped as Dump$^{neg}$ CD19$^{+}$ IgD$^{neg}$ CD38$^{neg}$ CD95$^{+}$ cells. Spike-specific B cell populations were identified as binding to recombinant SARS-CoV-2 Spike tetramers conjugated to PE or APC. The frequency of B cells dual stained with the two Spike tetramers is expressed as percentage ± standard deviation (STDEV).

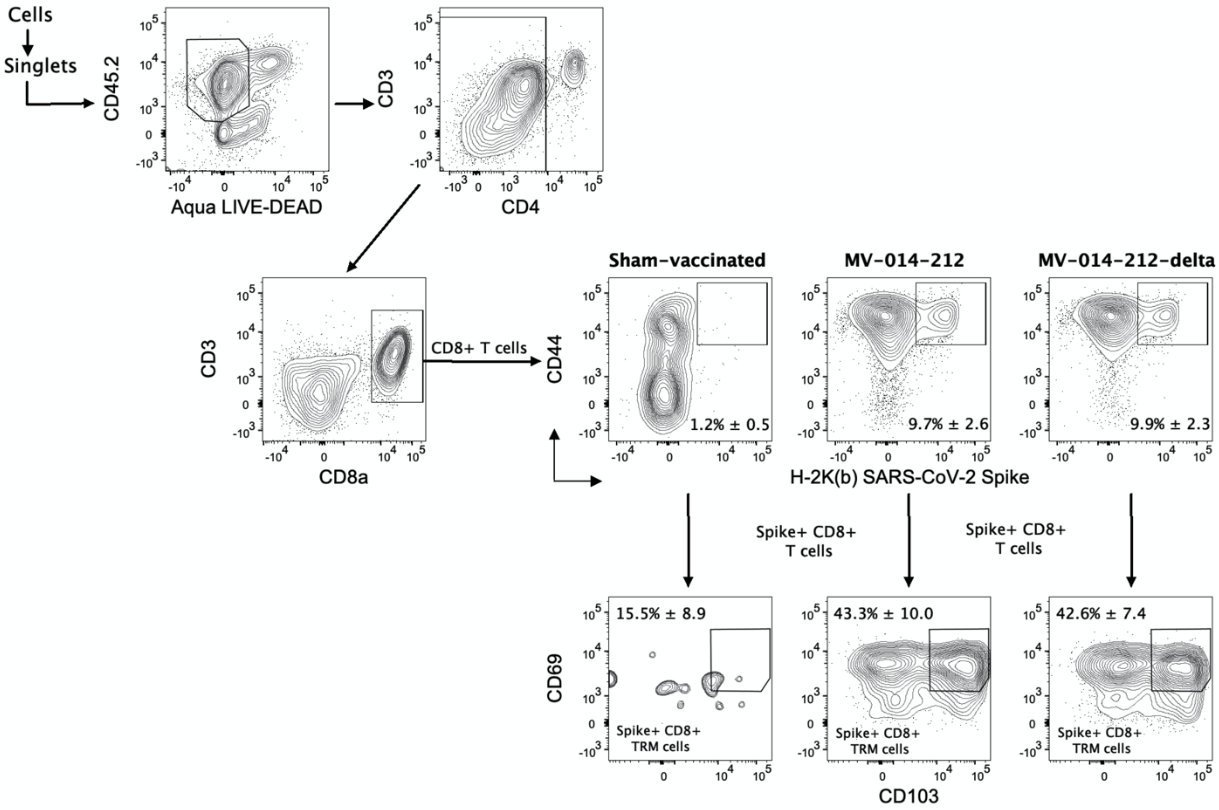


**Figure S3. Gating strategy for flow cytometric analysis of Spike-specific $CD8^+$ T cells in the lungs from vaccinated mice.** Sequential gating strategy for the identification and enumeration of total Spike-specific $CD8^+$ T cells and Spike-specific tissue-resident memory (TRM) $CD8^+$ T cells in the lungs of K18-hACE2 mice at day 10 post vaccination with Vero cell lysate (sham-vaccinated), MV-014-212 or MV-014-212-delta. $CD45.2^+$ live ($Aqua^{neg}$) cells were gated and total $CD8^+$ T cells were identified as $CD4^{neg}$ $CD3^+$ $CD8^+$ cells. Spike-specific $CD8^+$ T cells were identified with fluorochrome-labeled Class I MHC H-$2K^b$ SARS-CoV-2 Spike (VNFNFNGL)-BV421 tetramers. Spike-specific $CD8^+$ TRM cells were identified based on dual expression of $CD69^+$ and $CD103^+$. The frequency of each Spike-specific $CD8^+$ T cell population is expressed as percentage ± STDEV.

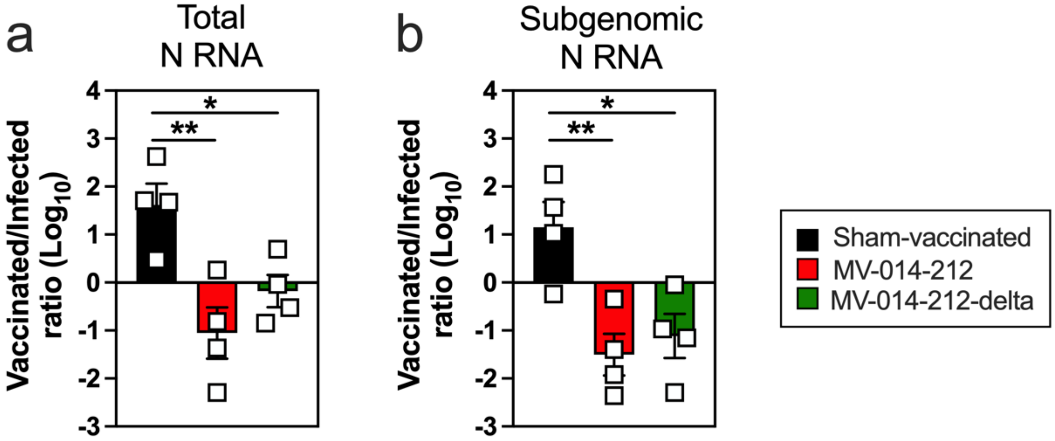


**Figure S4. Viral N transcripts in nasal swipe samples collected from infected animals and vaccinated cagemates.** Comparison of total (**a**) and subgenomic (**b**) viral N RNA levels from nasal swipes collected from experimentally-infected hamsters and their vaccinated (red and green) or sham-vaccinated (black) cagemates. Data are reported as the ratio of viral N RNA transcript levels (copies/ml) in nasal swipes from day 5 Delta SARS-CoV-2-infected animals to those collected from vaccinated/sham-vaccinated cagemates at day 5 post pairing. Animal pairs (n=4/group) are shown as symbols with the means of the ratios reported as bars. Data plotted as log-transformed ratios. *P <0.05, **P<0.01. P values determined by one-way ANOVA followed by Dunnett's multiple comparison test. Variability expressed as SE.
